# Runcer-Necromancer: A Method To Rescue Data From An Interrupted Run On MGISEQ-2000

**DOI:** 10.1101/2020.11.02.364588

**Authors:** Anna Pavlova, Vera Belova, Robert Afasizhev, Irina Bulusheva, Denis Rebrikov, Dmitriy Korostin

**Affiliations:** Center for Precision Genome Editing and Genetic Technologies for Biomedicine, Pirogov Medical University, 117997 Moscow, Russia

**Keywords:** MGISEQ-2000, DNBSEQ-G400, NGS, Paired-end sequencing, fastq merging

## Abstract

During the sequencing process, problems can occur with any device including the MGISEQ-2000 (DNBSEQ-G400) platform. We encountered a power outage that resulted in a temporary shutdown of a sequencer in the middle of the run. Since barcode reading in MGISEQ-2000 takes place at the end of the run, it was impossible to use non-demultiplexed raw data. We decided to completely use up the same cartridge with reagents and flow cell loaded with DNB and started a new run in a shortened custom mode. We figured out how the MGISEQ-2000 converts preliminary data in .cal format into .fastq files and wrote a script named “Runcer-Necromacer” for merging .fastq files based on the analysis of their headers (available online: https://github.com/genomecenter/runcer-necromancer). Read merging proved to be possible because the MGISEQ-2000 flow cell has a patterned structure and each DNB has invariable coordinates on it, regardless of its position on the flow cell stage. We demonstrated the correctness of data merging by comparing sample analysis results with previously obtained .fastq files for them. Thus, we confirmed that it is possible to restart the device and save both parts of the interrupted run.

## Introduction

At the end of 2017, Chinese company MGI Tech presented MGISEQ-2000 sequencing platform [1] promoting it as a device for large and medium scale genome sequencing. MGISEQ is specific in harnessing cPAS sequencing technology and using nanoballs (DNB) generated from circular molecules of DNA library by rolling circle replication [2]. MGISEQ is compatible with a wide range of reagents for sequencing in SE50, SE100, SE400, PE100, PE150, and PE200 modes. MGISEQ-2000 provides the quality of sequencing comparable with that of the Illumina platform [3–6].

The first MGISEQ-2000 sequencer in Russia was installed in our lab in February 2019, and we run it once a week in the paired-end 150 mode (PE150). According to our experience, one PE150 run usually takes 68 hours if one flow cell is used at a time. During one of these runs, about 23.00 on Saturday, there was a failure of the Moscow power grid leading to a 50-minute blackout of a whole district including Pirogov Medical University. UPS battery storage was sufficient only for 20 extra minutes, then the sequencer had been turned off until the power was restored. So the device with loaded reagents remained in a sleep mode for 35 h until Monday. Before the instrument was switched off, it had performed 138 full cycles of forward read sequencing (run 27). The specific feature of the MGISEQ-2000 sequencing program is that it reads a barcode at the end of a run after it completed sequencing of forward and reverse reads. So to a first approximation, the data obtained could not be demultiplexed as information on the barcodes was absent.

According to the MGI Tech [7] recommendations, after consulting with the MGI Tech service engineers, we were advised to dispose of the current tank with reagents as well as the flow cell and run the samples using new reagents. In the first place, it is linked to the high sensitivity of the MDA reagent to storage at +4 °C as it loses its activity very quickly. We decided to continue the run using the reagents that had been loaded for the weekend and try to restore the data. Finally, we managed to rescue the data using the software from ZebraCall [8] and our own script on C++.

## Materials and methods

### Sequencing

We prepared 3 pools of circularized libraries following the standard MGI Tech protocol [9]. Then we synthesized DNB, loaded a flow cell using the MGIDL-200H manual loader, prepared a sequencing cartridge from the MGISEQ-2000RS High-throughput Sequencing Set with User manual version: A2, and started sequencing on A-side in PE150 mode. Run 27 was aborted at the 139th cycle of the read-1 sequencing phase. After 35 h, we restarted the run (run 27_2) using the same sequencing cartridge and flow cell in a custom mode with the following parameters: read 1 for 12 cycles, read 2 for 151 cycles, Start phase: Sequencing (**Figure 1**). For summary reports generated by MGISEQ-2000 for lane 1 of runs 27 and 27_2 see File S1, S2.

**Figure 1.**
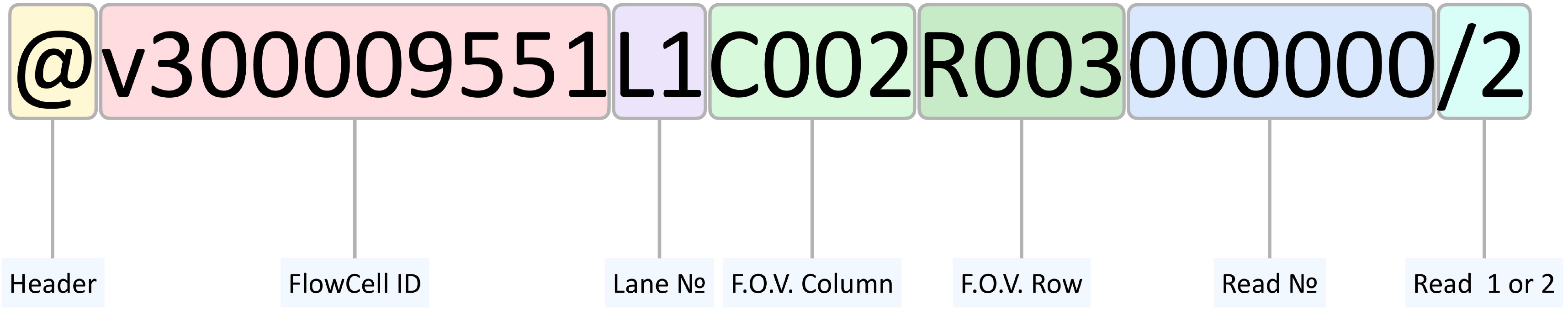
The screenshot of MGISEQ software in a custom mode with the settings used for restarting the run.

### fastq generation

The generation of .fastq files containing forward reads for the interrupted run was performed using ZebraCall v2 framework (C\:ZebraCallV2\client.exe – the pathway to software on MGISEQ-2000) which transforms intermediate .cal files into fastq format and demultiplexes them using barcodes.

The appropriate work of ZebraCall requires a .txt file with barcode sequences used for demultiplexing. We created an empty file ‘empty_barcode.txt’ so that the last 10 nucleotides from 13 nucleotides that were read earlier would not be recognized as barcodes by ZebraCall.

We used the following command (we provide an example for lane 1):

~~~
client.exe D:\Result\workspace\run_name\L01 139 6 72 -B
C:\ZebraCallV2\empty_barcode.txt -N run_name -U 1 -F
~~~

It contains the options:

the access to the folder with .cal files
run_name — the name of a run
139 – the number of completed sequencing cycles
6 72 – the number of fields of view counted horizontally and vertically for a corresponding lane
-B – a path to the file with barcodes
-U – the number of a lane
-F – fastq generation without generation of flow cell images

As a result, for each lane, we generated files ‘run_name_L0N_read.fq.gz’ where N is a lane number. Such file contained a read name and a sequence of 138 nucleotides long.

### Fastq merging

MGISEQ-2000 employs a patterned flow cell, so each DNB in a cell has unique coordinates at X and Y axes which do not depend on flow cell localization in a device and are not changed if the flow cell is displaced. When the power of the sequencer was off, the vacuum pump was switched off as well. The coordinates of each read were saved in a header of a .fastq file (**Figure 2**). This allowed us to integrate the data on forward reads obtained before and after the instrument was off.

**Figure 2.**
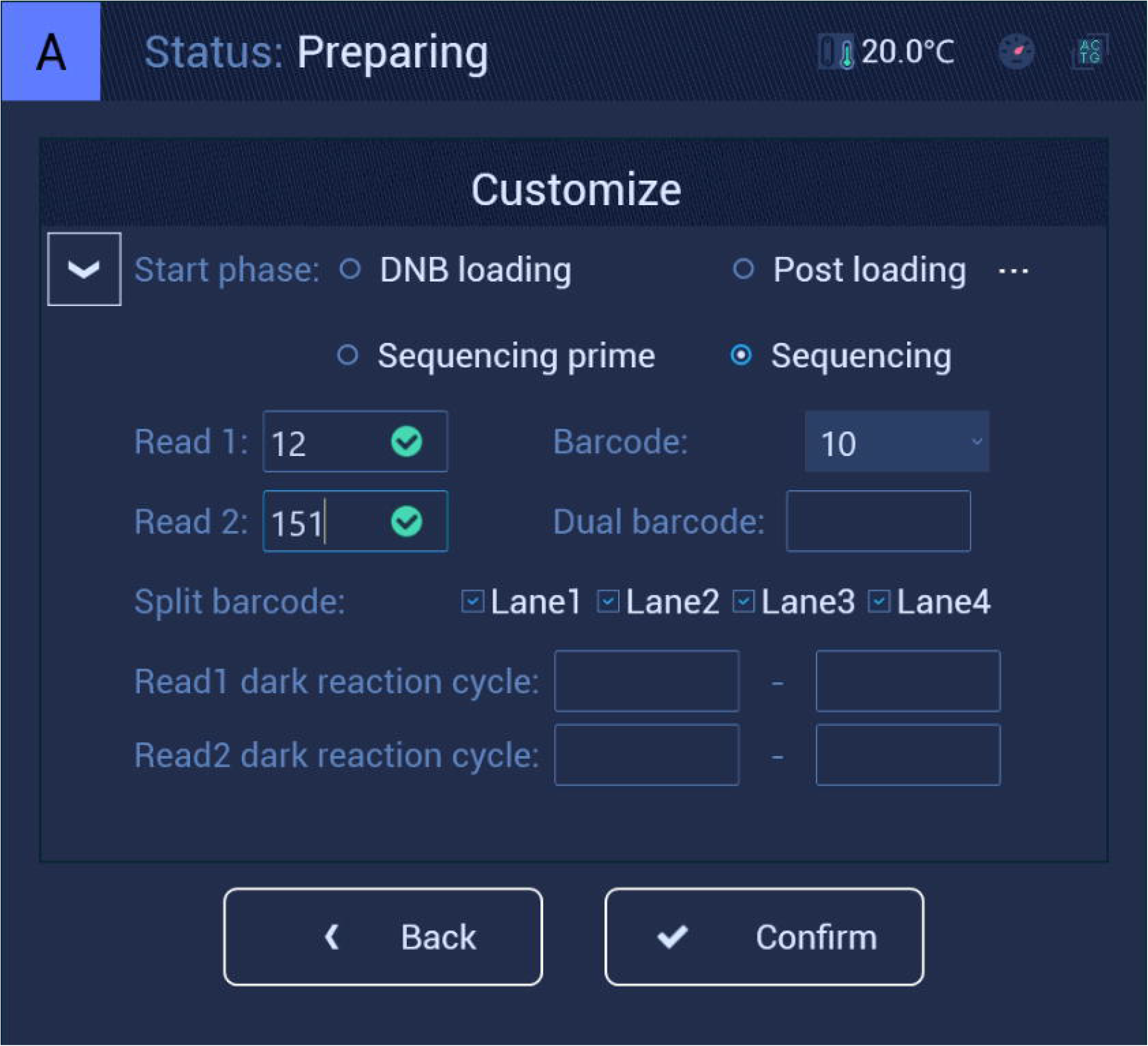
The structure of a read header in an MGISEQ-2000 .fastq file.

**Figure 3.**
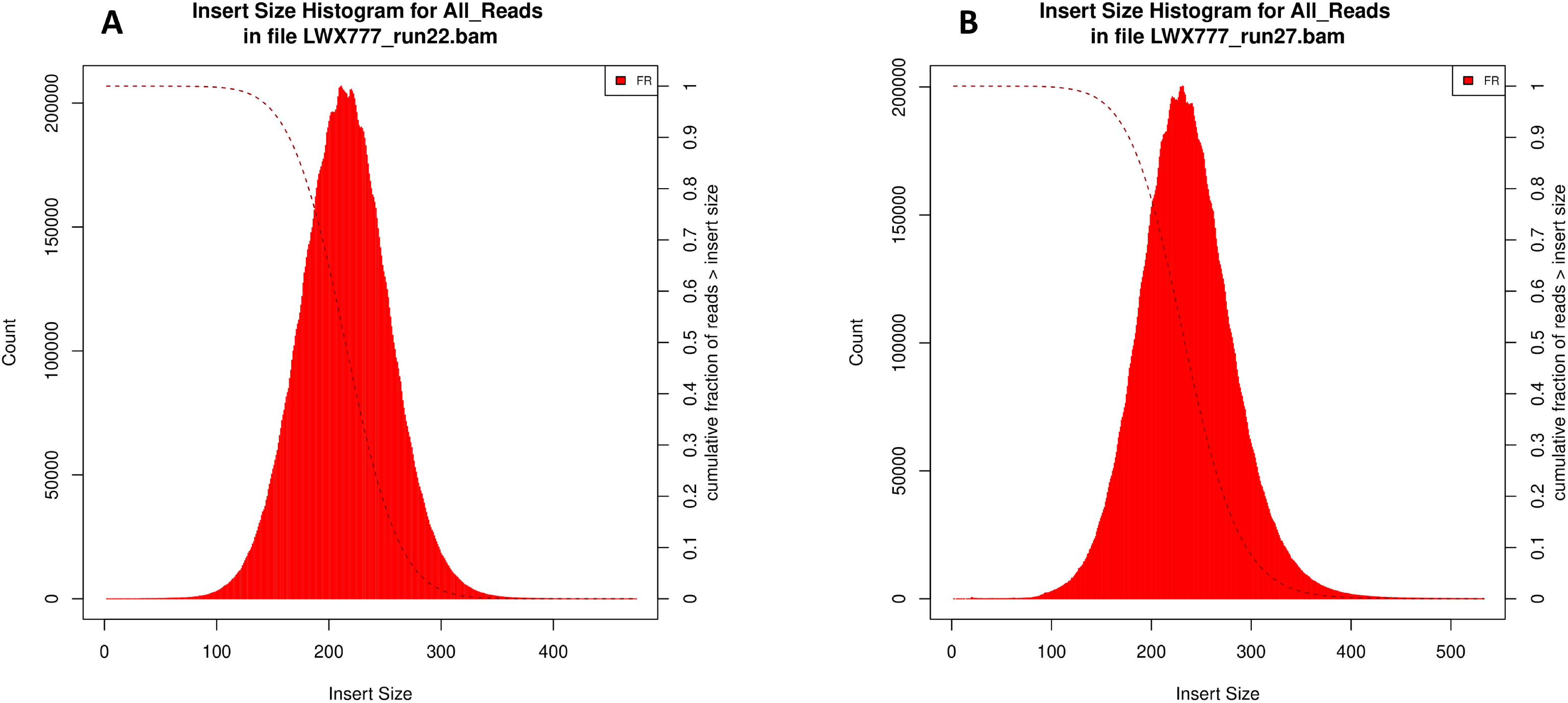
Q30 histograms for runs 27 (A) and 27_2 (B) The X-axis represents the number of sequencing cycles, the Y-axis represents the ratio of the data with the quality no less than Q30 (%). Blue arrows in histogram B indicate the cycles which reverse reading and barcode reading started.

As a read number being used for forward and reverse reading is unique, we managed to combine the 138-nucleotide sequences obtained during the first run with the nucleotide sequences obtained during the second run based on the information on F.O.V Column, F.O.V Row, and read numbers. To achieve this, we created a C++ script which can be accessed at GitHub [10]. The instruction for script running can be found in the file README.md in the repository.

## Results

The most important parameter for sequencing quality is the ratio of the data with the quality level of no less than Q30. The Q30 value and other quality metrics did not decrease dramatically in spite of a 35hour stand by (**Figure 2**, **Table 1)**.

**Table 1.** Metrics of 27 and 27_2 run for lane 1 from Summary report (File S1, S2)

| **Run** | **27** | **27_2** | **Drop** |
| --- | --- | --- | --- |
| Loaded DNBs (from the first base<br>report) | 1169312 | 1131610 | 3% |
| Chip productivity, % | 76.82 | 68.18 | 11% |
| Q30, % | 90 | 90.96 | 0% |
| Total reads per lane, M | 467.3 | 397.67 | 15% |

This implies that storing a loaded cartridge for 35 h leads to its decline, however, it can be still used for sequencing.

To check if merging reads from different runs was correct, we compared 7 samples of whole-exome sequencing from runs 27 and 27_2 with the data from the same samples obtained in a run 22. We used the distribution of the size of an insert between left and right reads and the ratio of reads having the insert size exceeding 1000 nucleotides as control metrics. If read merging had been performed with errors, the portion of the reads mapped to various genome regions would have significantly increased. See **Figure 4** for the distribution of an insert size for sample LWX777 from the control group. The ratio of reads from sample LWX777 with the insert size exceeding 1000 nucleotides was 0.003% in case of the data combined from the different runs, while it was 0.005% in case of the previous sequencing without read integration. The obtained data imply that read merging was correct.

**Figure 4.**
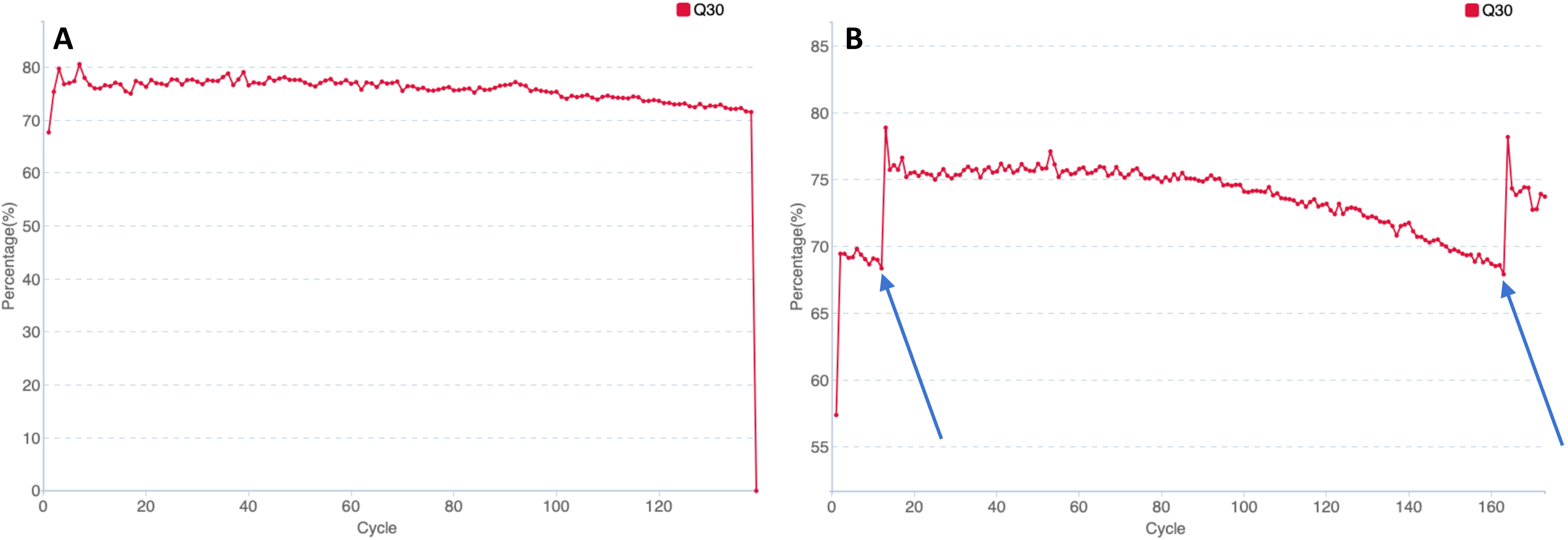
The distribution of inserts for sample LWX777 from runs 22 (A) and 27+27_2 (B) The X-axis represents the number of reads, the Y-axis represents an insert size. The diagrams were obtained using Picard CollectIsertSizeMetrics.

## Conclusion

It is possible to use a sequence cartridge after 35-hour storage at +4°C, although the quality of the obtained data is reduced.

Merging sequencing data can be successfully performed if the information about the localization in flow cells is saved in a read header.

## Authors’ contributions

AP performed custom fastq merging and proved its correctness. VB prepared samples for sequencing and started run. RA participated in data analysis. IB converted raw data with ZebraCall. DR participated in study design. DK conceived of the study and supervised it and drafted the manuscript. All authors read and approved the final manuscript.

## Supplementary material

**File S1 Summary report for run 27 lane 1 from MGISEQ-2000**

**File S2 Summary report for run 27_2 lane 1 from MGISEQ-2000**

